# Visual perturbation of balance suggests impaired neuromuscular stability but intact visuo-motor control in Parkinson’s disease

**DOI:** 10.1101/2021.07.05.451110

**Authors:** David Engel, Justus Student, Jakob C.B. Schwenk, Adam P. Morris, Josefine Waldthaler, Lars Timmermann, Frank Bremmer

**Author notes:** Corresponding Author: David Engel, Dept. Neurophysics, Philipps-Universität Marburg, Karl-von-Frisch-Str. 8a, 35043 Marburg, Germany.

## Abstract

Postural instability marks one of the most disabling features of Parkinson’s disease (PD), but only reveals itself after affected brain areas have already been significantly damaged. Thus, there is a need to detect deviations in balance and postural control before visible symptoms occur. In this study, we visually perturbed balance in the anterior-posterior direction using sinusoidal oscillations of a moving room in virtual reality at different frequencies. We tested three groups: individuals with PD under dopaminergic medication, an age-matched control group, and a group of young healthy adults. We tracked their centre of pressure and their full-body motion. We investigated sway amplitudes and applied newly introduced phase-locking analyses to investigate responses across participants’ bodies. Patients exhibited significantly higher sway amplitudes as compared to the control subjects. However, their sway was phase-locked to the visual motion like that of age-matched and young healthy adults. Furthermore, all groups successfully compensated for the visual perturbation by – most likely reflexively - phase-locking their sway to the stimulus. As frequency of the perturbation increased, distribution of phase-locking (PL) across the body revealed a shift of the highest PL-values from the upper body towards the hip-region for young healthy adults, which could not be observed in patients and elderly healthy adults. Our findings suggest an impaired neuromuscular stability, but intact visuomotor processing in early stages of PD, while less flexibility to adapt postural strategy to different perturbations revealed to be an effect of age rather than disease.

**New & Noteworthy:** A better understanding of visuomotor control in Parkinson’s disease (PD) potentially serves as a tool for earlier diagnosis, which is crucial for improving patient’s quality of life. In our study, we assess body sway responses to visual perturbations of the balance control system in patients with early-to-mid stage PD, using motion tracking along with recently established phase-locking techniques. Our findings suggest patients at this stage to have an impaired muscular stability but intact visuomotor control.

## Introduction

Second only to Alzheimer’s, Parkinson’s disease (PD) is one of the most prevalent neurodegenerative diseases and – accompanied by our aging demography - has an ever-growing impact on modern society (Elbaz et al., 2016). The exact causes of PD are still unknown. However, there is agreement that the basal ganglia constitute the main brain area affected (Abbruzzese & Berardelli, 2003), where dying of dopaminergic neurons in the substantia nigra (SN) leads to reduced function (Elbaz et al., 2016; Chen et al., 2016). The basal ganglia are involved in processing and integrating (multi-)sensory information, especially in determining subsequent motor output (Bolam et al., 2002; Abbruzzese & Berardelli, 2003; Nagy et al., 2006). As a consequence, main symptoms of PD include bradykinesia, tremor, as well as rigidity and postural instability (Bloem, 1992; Hwang et al., 2016; Feller et al., 2019). Impairment of balance and posture is generally seen as one of the most disabling features of PD (Bloem, 1992; Grimbergen et al., 2009; Hwang et al., 2016). It is followed by an increased risk of falls, which considerably reduces the quality of life (Koller et al., 1989; Horak, 2006; Benatru et al., 2008, Hwang et al., 2016; Doná et al, 2016).

Crucially, motor symptoms only start to occur after about half of the cells in the SN have deceased (Fearnley & Lees, 1991). This includes postural instability, which most commonly emerges several years after the onset of motor symptoms, and is, thus, a symptom of advanced PD (Koller et al., 1989; Bloem, 1992; Hwang et al., 2016). Nevertheless, this late onset mostly accounts for those symptoms of postural instability which are apparent to a human observer and reveal themselves through simple clinical tests (Landers et al., 2008). Impairments potentially occur significantly earlier in the progression of the disease and might be detectable in more nuanced and sophisticated measures of posture and sway. In addition, postural stability is thought to be comprised of two sub-systems, a ‘passive’ neuromuscular system and an ‘active’ sensorimotor system, which might be affected differently by PD (Bloem, 1992; Chen et al., 2016). Since there is still no cure for the disease, early diagnosis is the most effective tool against PD, as it allows for counter measures to be taken at early stages, facilitating better treatment and a sustainable quality of life for those affected. Thus, there is a need for biomarkers to detect the disease closer to its onset, before easily visible (motor) symptoms occur.

Due to the ever-changing sensory inputs in every-day life, the human balance control system is constantly being supplied with new information to which it must respond. This typically results in a constant sway of our body around its point of equilibrium (Horak & MacPherson, 1996; Schoneburg et al., 2013). The three main sensory inputs we use to maintain balance are vision, graviception (vestibular input) and proprioception (Bronstein et al., 1990; Horak & MacPherson, 1996; Azulay et al., 2002; Peterka & Loughlin, 2004; Horak, 2006). Investigation of this body sway, especially how it adapts under perturbations, gives insight into the underlying sensorimotor system and its processing (Lee & Lishman, 1975; Peterka, 2002; Musolino et al., 2006). It has been proposed that a major aspect of postural instability in PD is an impaired perception of movement rather than execution of movement (Richards et al., 1993; Hwang et al., 2016; Halperin et al., 2020). This includes perception of self-motion (Yakubovich et al., 2020). In this context, PD patients seem to show a higher dependence on visual information for motor and posture control (Cooke et al., 1978; Bronstein et al., 1990; Azulay et al., 2002; Weil et al., 2016; Bronstein, 2019), which might be attributable to proprioceptive deficits (Abbruzzese & Bernardelli, 2003; Keijsers et al., 2005; Jacobs & Horak, 2006; Benatru et al., 2008). However, this increased dependence on vision has also been observed in older healthy adults and might be an effect of age, rather than disease (Wade et al., 1995; Toledo et al., 2014). Nevertheless, visual information that perturbs the postural control system and particular nuances in the reaction of said system might be indicative of early visuomotor impairments within the progress of PD.

A well-established procedure for visually perturbing the balance control system is the ‘moving-room’ paradigm, in which the visual environment is moved around the observer, eliciting the illusion of self-motion and inducing *visually evoked postural responses* (VEPR, Lestienne et al., 1977; Bronstein et al., 1990; Schöner, 1991). Since they are easy to implement and have strong analytical benefits, periodic perturbations in the form of sinusoidal oscillations of the visual surrounding have proven to be a reliable tool for the investigation of VEPR (Lee & Lishman, 1975; Schöner, 1991; Scholz et al., 2012; Hanssens et al., 2013; Cruz et al., 2018; Engel et al., 2020). Experiments of this kind have been performed with PD patients before (Bronstein et al., 1990; Hwang et al., 2016), most notably in the recent work of Cruz and colleagues (2018, 2020, 2021), who used oscillatory visual stimuli at different frequencies. In these experiments, using one kinematic measure (an optical motion tracker placed on the back), PD patients showed corrective responses to the visually moving room similar to age-matched controls.

Body sway is commonly measured by tracking the foot centre of pressure (COP) with a force plate, which might be supported by additional motion tracking of the upper body (Lestienne et al., 1977; Winter, 1995; Jeka et al., 1998; Jacobs & Horak, 2006; Scholz et al., 2012; Boonstra et al., 2016; Engel et al., 2020, 2021). In recent years, measuring devices adopted from the video game industry are increasingly being established for research purposes. In particular, the Nintendo Wii Balance Board and the Microsoft Kinect v2 have extensively been validated (Dehbandi et al., 2017; Clark et al., 2018). Combined with head-mounted virtual reality headsets which allow for convenient presentation of immersive visual stimuli, these devices provide cost-effective and mobile means to assess postural control (Garner & D’Zmura, 2020; Engel et al., 2021).

Prevalent methods for analysing the measures of body sway mentioned above include general sway magnitude, often expressed as root mean square (Barela et al., 2009; Cruz et al., 2018; Feller et al., 2019), as well as frequency spectrograms to gain insight into the frequency content of the signals (Loughlin & Redfern, 2001; Creath et al., 2005; Musolino et al., 2006; Laurens et al., 2010; Engel et al., 2020). Frequency analyses are particularly effective when the perturbation is periodic, as one can obtain the balance system’s dynamic response to the perturbation (Schöner, 1991; Musolino et al., 2006; Scholz et al., 2012; Cruz et al., 2018, 2020, 2021; Engel et al., 2020, 2021). However, investigation of both body sway magnitude and spectral analyses based on frequency power have revealed high inter-subject variability and thus often lead to conflicting results, making it difficult to obtain comparable responses to the stimuli used in experiments (Kay & Warren, 2001; Sparto et al., 2004; Chastan et al., 2008; Feller et al., 2019; Cruz et al., 2018, 2020, 2021). To overcome these obstacles, phase coherence analyses independent of frequency power have recently proven to be a powerful instrument to investigate responses to oscillatory stimuli (Engel et al., 2020). This includes obtaining the phase-locking value (PLV), which has been adopted from EEG-studies and evaluates phase consistency of a signal over time and across trials (Lachaux et al., 1999). Recently, we applied PLV analyses to full-body motion data in response to periodical visual stimulation, which allowed us to observe a shift in coordination strategy as a function of stimulus frequency in healthy adults (Engel et al., 2021).

Taken together, current evidence on PD suggests an impaired balance response to visual stimuli, which might be evident in subtle posturographic measures. As recently established phase analyses yielded well-nuanced and reliable responses to oscillatory visual drives in healthy adults, it is intriguing to investigate how patients suffering from PD, based on kinematic data of their entire body, phase-lock to these stimuli. To incorporate possible aging-effects, it is vital to also include young healthy adults in the experiments. Thus, in this study, we used a low-cost and mobile experimental setup to implement a sinusoidal moving room paradigm at different frequencies in virtual reality that has not been previously applied to study PD. We assessed VEPR of patients with PD and two groups of participants without history of neurological impairments (i.e., age-matched and young adults) by tracking their COP and 25 body segments and analysing the signals in terms of sway magnitude and PLV. We hypothesized (1) that PD patients show greater sway magnitude at all frequencies of stimulation due to their postural instability. Moreover (2), that due to their increased visual dependence, they are more susceptible to the visual stimulus and thus exhibit exaggerated phase-locking. Thirdly (3), based on the common symptom of rigidity, that PD patients use less flexible coordination strategies across their bodies in response to the visual movement in comparison to both groups of healthy adults.

## Materials and Methods

### 2.1 Participants

The group of participants consisted of three cohorts. We collected data of 15 healthy young subjects (22–29 years, mean = 24.4 ± 2.2 years; 11 males, 4 females), 12 elderly healthy subjects as age-matched controls (48-70 years, mean = 58.3 ± 6.9 years; 7 males, 5 females) and 18 patients diagnosed with PD according to the Movement Disorders Society clinical diagnostic criteria for Parkinson’s disease (Postuma et al., 2015; 42-76 years, mean = 58.1 ± 8.9 years; 16 males, 2 females; Hoehn and Yahr Scales 1-3; all “on” dopaminergic medication [levodopa equivalent dosage(LED): 651.63 ± 529.97]). They will be referred to as YOUNG (YG), CONTROL (CT) and PD, respectively. Participants of all three groups had normal or corrected to normal vision and no known orthopaedic, musculoskeletal or neurological impairments. All subjects gave written informed consent prior to the experiment, including with regard to the storage and processing of their data. Data was collected at two locations, while all patients were recruited from University Hospital Marburg. Experimental procedures performed at Monash University, Melbourne, Australia were approved by the Monash University Human Research Ethics Committee (17956) and were conducted in accordance with the National Statement on Ethical Conduct in Human Research. Experimental procedures performed at the University of Marburg, Marburg, Germany were approved by the Ethics Committee of the Psychology Department, University of Marburg. Research including PD patients was approved by study 77/19 of the Ethics Committee of the Faculty of Medicine, University of Marburg. All research was conducted in accordance with the Declaration of Helsinki.

### 2.2. Experimental Setup and Stimulus

Our setup and stimulus have been established and used in a previous study (Engel et al., 2021). Visual stimuli were presented through a head mounted display (HTC Vive, HTC, New Taipei City, Taiwan) with a frame rate of 90 Hz. The headset’s field of view spanned over the central 110° in both vertical and horizontal directions. Subjects’ COP was tracked using a Nintendo Wii Balance Board (WBB, Nintendo, Kyoto, Japan). We additionally performed full body motion tracking using a Microsoft Kinect v2 (Microsoft, Redmond, WA, USA) which provides tracking of 25 ‘body joints’ based on an internal skeleton model in 3-D (Dehbandi et al., 2017). The Kinect was located at a distance of 210 cm in front of the WBB, at a height of 140 cm at both recording sites, Melbourne and Marburg, respectively. Subjects wore no shoes and were asked to stand relaxed with their feet about hip width apart and parallel on the WBB. They were instructed to let their arms dangle at the side of their body without effort and to maintain their gaze straight ahead. Throughout the experiment, participants wore a harness which was connected to a beam at the ceiling. The harness was arranged in such a way that it ensured the participants’ safety but did not provide any lift during trials. We used a custom-built 3-D virtual environment based on the Python pyopenvr framework in *OpenGL* to create the visual stimulus. It consisted of a 3-D tunnel made up of black spheres (GLPoint_size = 1000 at zero distance, density = 50 per unit cube). The positions of the spheres were randomly generated along the walls of the tunnel at each trial. Their size scaled inversely with distance (the closer the larger). Subjects stood in a grey isotropic infinite space, in which the tunnel was world-fixed, its origin placed at their eyes. It stretched into the anterior-posterior direction. We arranged the position of the WBB and the virtual world in such a way that subjects were facing the Kinect. The length of the entire tunnel was set to 50 m. The radial centre was individually adjusted to each participant’s eye level. This gave them the impression of standing on the ground of the tunnel. We implemented a fixation dot (GLPoint_size = 5) at participant’s eye level and a distance of 24 m, close to the end of the tunnel.

### 2.3. Experimental Paradigm

The paradigm included three oscillatory movement conditions and a baseline condition, during which the tunnel remained static. In each movement condition, the tunnel first remained static for 5-8 seconds (randomized) and then oscillated along the sagittal plane (anterior-posterior) for 30 seconds at distinct frequencies of 0.2 Hz, 0.8 Hz and 1.2 Hz with a randomized starting phase. The movement of the tunnel was followed by another 5 s when it remained static to provide the subjects with relaxation time (Figure 1). We scaled the amplitude of each oscillation with the inverse of its frequency to maintain a constant speed of the optic flow and to prevent velocity effects (Dokka et al., 2009; Hanssens et al., 2013). The default amplitude was set to an equivalent of 1 cm. This made the tunnel oscillate just above visual detection threshold at the highest frequency.

**Fig. 1.**
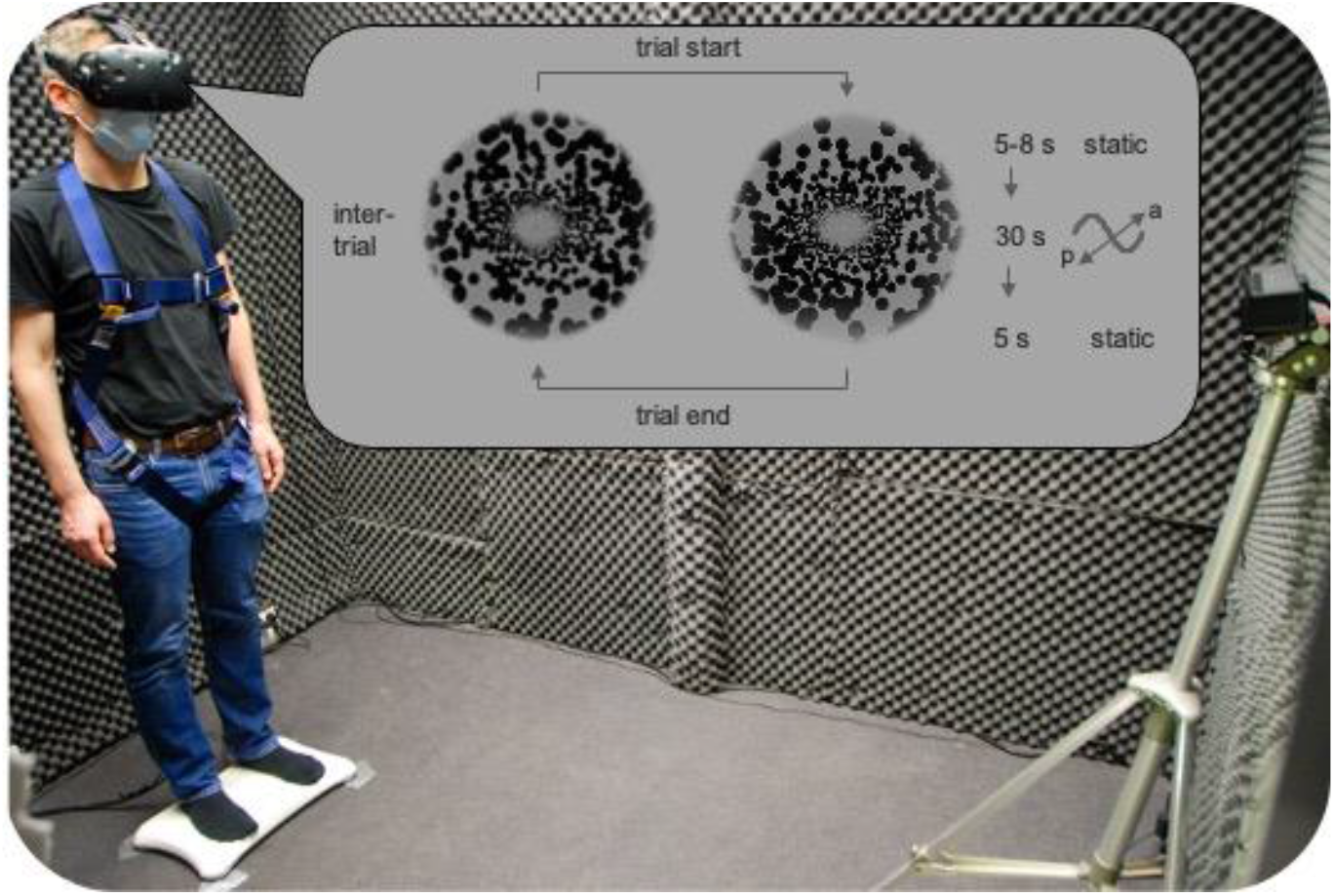
Experimental setup. Participants wore a virtual reality headset for visual stimulation. They stood on a Wii Balance Board to measure their COP and their body motion was captured using a Kinect markerless motion tracking camera. The headset simulated them standing inside a tunnel. In each motion trial, after a static period of 5-8 s, the tunnel oscillated in the anterior-posterior direction for 30 s at one of three frequencies (0.2 Hz, 0.8 Hz, 1.2 Hz), after which it remained static again for 5 s. For details see 2.2.

Each subject performed 10 trials per condition in pseudorandom order. The beginning of each trial was indicated by the fixation dot changing its colour to white. Following this cue, participants stood as instructed and fixated. To ensure that subjects kept fixating and thus keep their peripheral visual field stable (Horiuchi et al., 2017; Raffi & Piras, 2019), we implemented a counting task which included transient colour flips of the fixation dot. Subjects had to count the number of flips and report the total number at the end of each trial.

Between trials, the tunnel remained static, and subjects could stretch and rest for as long as they needed. These resting periods were indicated to the subjects by a red fixation dot and the following trial was started by the experimenter at their verbal command. To prevent fatigue, trials were distributed across a minimum of three blocks, between which we arranged for breaks where subjects were able to leave the setup and rest. All trials were recorded in a single session on the same day.

### 2.4. Data analysis and statistics

We used custom-made Python programs for initial raw data recording and storage. All subsequent data processing and analyses were performed in MATLAB (The MathWorks, Inc., Natick, USA). Since both the WBB and the Kinect provide rather inconsistent sampling rates (around 100 Hz and 30 Hz, respectively), data collected from both devices was resampled at 50 Hz using a Gaussian moving average filter with a symmetric window (sigma = 1/60 s). Apart from this, no further filtering was applied to the raw data. Trial data was cut to the 30 s where the tunnel was moving (or remained stationary). For all subsequent analyses, we extracted the respective time-courses of the COP and the 3-D body segments corresponding to the anterior-posterior (A-P) direction.

To gain insight into the general degree of body sway across the different conditions, we used mean sway amplitude (MSA) which has been implemented before (Barela et al., 2009; Cruz et al., 2018). For this purpose, prior to calculating the standard deviation of the time courses, we subtracted a first-order polynomial from the raw time courses using the *detrend* function in MATLAB.

To investigate phase coupling of the bodily responses to the stimuli, the single trial time courses of the COP and the body segments were z-transformed and underwent a continuous complex wavelet decomposition, using the *cwt* function in MATLAB with generalized Morse wavelets (gamma = 3, 10 voices per octave). The resulting complex wavelet spectra were phase-corrected for the randomized phase onset of each trial. To avoid edge artifacts, only wavelet coefficients that were inside the cone of influence were considered. Wavelet frequency limits were chosen separately for each condition in order to match the discrete frequency bins of the resulting spectra with the exact stimulus frequencies.

Based on the acquired wavelet spectra, we used recently established phase-locking analyses adopted from EEG-studies (Lachaux et al., 1999; Engel et al., 2021) to analyse phase behaviour of the recorded COP and body segment responses. For this purpose, we normalized the wavelet coefficients to have a magnitude of one and subtracted the phase of a continuous sinewave at each respective frequency band (Eq. 1).

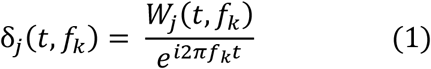

*δ_j_*: normalized complex coefficients with relative phase to the stimulus at *j*-th trial and *k*-th condition, *W_j_*: normalized wavelet coefficients, *f_k_*: frequency at condition k, *t*: time point

At the three stimulus-frequencies (conditions), these sinewaves resembled the time-course of the stimulus. The performed subtraction hence represents the phase difference between the responses and the visual stimuli. Out of these transformed complex wavelet coefficients, we calculated the phase-locking values (PLV) by taking the average across time points for each trial and subsequently the average across trials for each subject (Eq. 2).

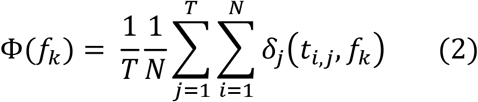

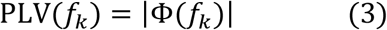

*PLV*: Phase-Locking Value, *N*: number of time points, *T*: number of trials

The absolute of those values then constituted the final PLVs (Eq. 3). These values represent the phase-locking to the stimulus across time and trials at each frequency band and range from 0 (entirely inconsistent relative phase) to 1 (entirely consistent relative phase). Thus, the phase-locking is independent of frequency power and reflects consistency of phase between the stimulus and the bodily response over time and across trials.

Due to unequal group sizes and high inter-subject variability, normality and homogeneity of variance could not be assumed. Therefore, we applied non-parametric testing on ranked data to our results. On each obtained measure, to test if group results (PD, CT, YG) came from different populations, we performed an independent samples Kruskal-Wallis test with follow-up pairwise comparisons with adjusted p-values. To test for influence of the stimulus frequency on the obtained measures for each group, we performed related-samples Friedman’s two-way ANOVA, also including follow-up pairwise testing with adjusted p-values. We considered 95% confidence intervals (p<0.05) to reject the null hypothesis. For all follow-up tests, if significant, effect sizes were calculated by dividing the respective standardized test statistic by the square root of total participants in each case. Statistical analyses were performed in SPSS (IBM, Armonk, NY, USA).

Body plots (Fig. 3 & 6) were created using the fieldtrip toolbox (https://fieldtriptoolbox.org) along with a custom body scheme (modified from: *MenschSDermatome* by Uwe Thormann (Licensed under CC BY-SA 3.0). *BrewerColormaps* (Cobeldick, 2020) were used for visual representation of MSA and PLV.

## 3. Results

### 3.1. Mean Sway Amplitude (MSA)

Figure 2 shows the average mean sway amplitude (MSA) of the COP for all groups across all conditions. PD patients exhibited significantly more sway in their COP than the age-matched healthy controls and the healthy young subjects, while the latter showed the least sway in all cases. There was a significant effect of the group on average MSA in all four conditions (Static: H(2)=19.31, p<.001; 0.2 Hz: H(2)=13.62, p=.01; 0.8 Hz: H(2)=18.25, p<.001; 1.2 Hz: H(2)=12.81, p=.002). For the static condition, follow-up pairwise comparison with adjusted p-values revealed significant differences between PD and both other groups (PD-YG: p<.001, r =.74; PD-CT: p=.019, r=.50). Stimulation at 0.2 Hz revealed a significant difference between the PD and the YG group (p=.001, r=.63). Stimulation at 0.8 Hz again led to significant differences between PD and both other groups (PD-YG: p<.001, r=.73; PD-CT: p=.029, r=.47). Stimulation at 1.2 Hz led to a significant difference between PD and YG (p=.001. r=.61).

**Fig. 2.**
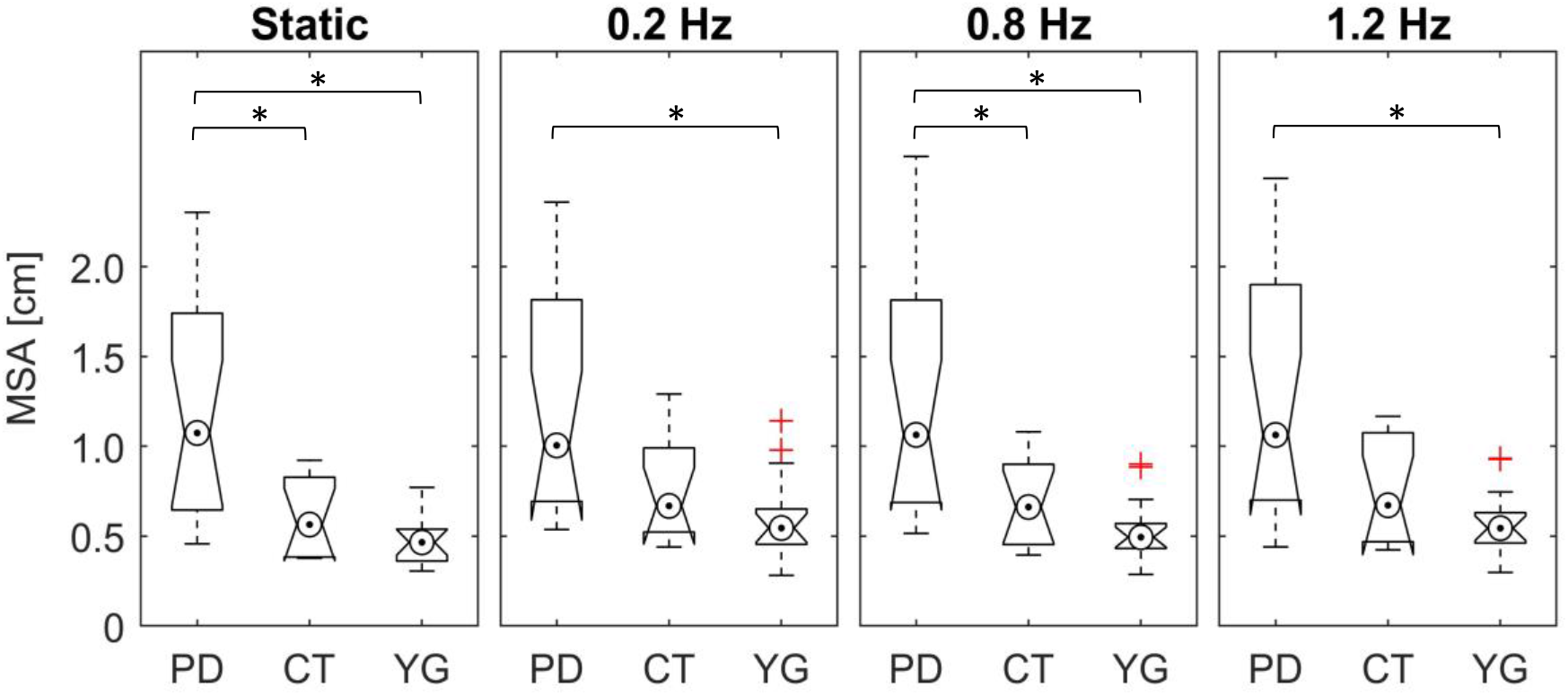
Boxplots and whiskers of COP mean sway amplitude (MSA) for the three tested groups across all conditions (Static = no tunnel motion) in A-P direction. Columns from left to right represent stimulus condition. Within columns, boxplots represent data from each respective group of PD patients (PD), age-matched controls (CT) and young healthy adults (YG). Crosses indicate outliers.

**Fig. 3.**
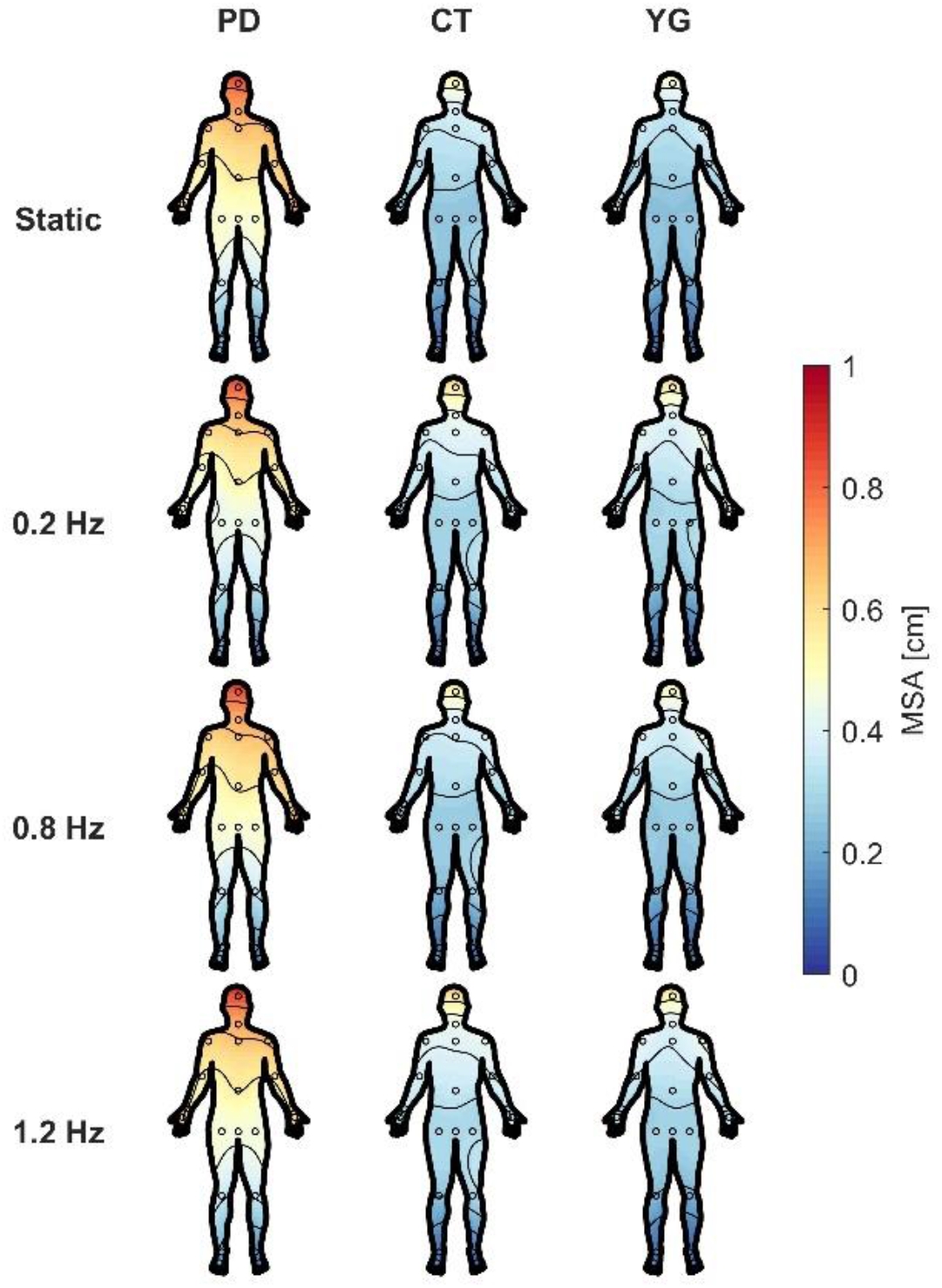
A-P mean sway amplitude (MSA) of the 25 body segments obtained by the Kinect. Color-coding represents group average of MSA for each body segment. The circles represent single body segments. Columns from left to right show groups (PD, CT, YG). Rows from top to bottom show different frequencies of the simulated tunnel movement (Static, 0.2 Hz, 0.8 Hz, 1.2 Hz).

For PD patients, motion of the tunnel at any frequency had no significant effect on overall sway of the COP in A-P direction (χ^2^(3)=6.87, p=.076). Among the age-matched controls (CT) however, there was a significant influence of the condition on MSA (χ^2^(3)=22.5, p<.001). Here, the static tunnel led to significantly lower MSA in follow-up pairwise comparison to all conditions where the tunnel was oscillating (0.2 Hz: p<.001, r=1.32; 0.8 Hz: p=.027, r=.82; 1.2 Hz: p=.005, r=.96). For the young healthy adults, there was also a significant effect of tunnel movement on their MSA (χ^2^(3)=14.76, p=.002). Follow-up pairwise comparison led to significant differences between the static and the 0.2 Hz condition (p=.002, r=.91) as well as between the static and the 1.2 Hz condition (p=.018, r =.77).

Average MSA of the body segments in A-P direction as measured by the Kinect can be seen in Figure 3. Analogous to the COP data, PD patients exhibited substantially more sway than both elderly and young healthy adults. In all groups, sway was most prominent at the head and decreased towards the feet. As was the case with the COP responses, the movement condition of the tunnel had little or no effect on body sway magnitude and did not affect distribution of sway across the body.

### 3.2. Phase-Locking Value (PLV)

To gain insight into phase-locking of participants’ responses to the stimulus, we first calculated the group average PLV for the COP data in A-P direction. The obtained PLV spectra are displayed in Figure 4. All groups showed a strong phase-coupling of their COP at the stimulus frequency, which is represented in the high and distinct peaks in each spectrum. As can be seen in the corresponding baseline spectra, PLVs did not show any peaks when the tunnel remained static. Since all groups revealed to have strong responses across conditions, we evaluated the height of the peaks for potential differences. Average peak heights for each frequency at which the tunnel was oscillating can be seen in Figure 5. There was only a significant effect of the group on COP PLV when the tunnel oscillated at 1.2 Hz (0.2 Hz: H(2)=1.10, p=.577; 0.8 Hz: H(2)=.27, p=.874; 1.2 Hz: H(2)=8.326, p=.016). Here, the follow-up pairwise comparison led to a significant difference between PD and CT (p=.012, r=.53). However, within each group, the stimulus affected the PLV magnitude significantly. For the PD patients (χ^2^(2)=18.78, p<.001), follow-up pairwise comparison revealed significant differences between 0.2 Hz and 0.8 Hz (p=.001, r=.86) and between 0.2 Hz and 1.2 Hz (p<.001,r=.90). For the age-matched controls (χ^2^(2)=13.50, p=.001), there was a significant effect between the lowest frequency of 0.2 Hz and the highest frequency of 1.2 Hz (p=.001, r=1.06). For the young healthy adults, (χ^2^(2)=11.20, p=.004), there was also a significant difference in PLV between 0.2 Hz and 1.2 Hz (p=.003, r=.62).

**Fig. 4.**
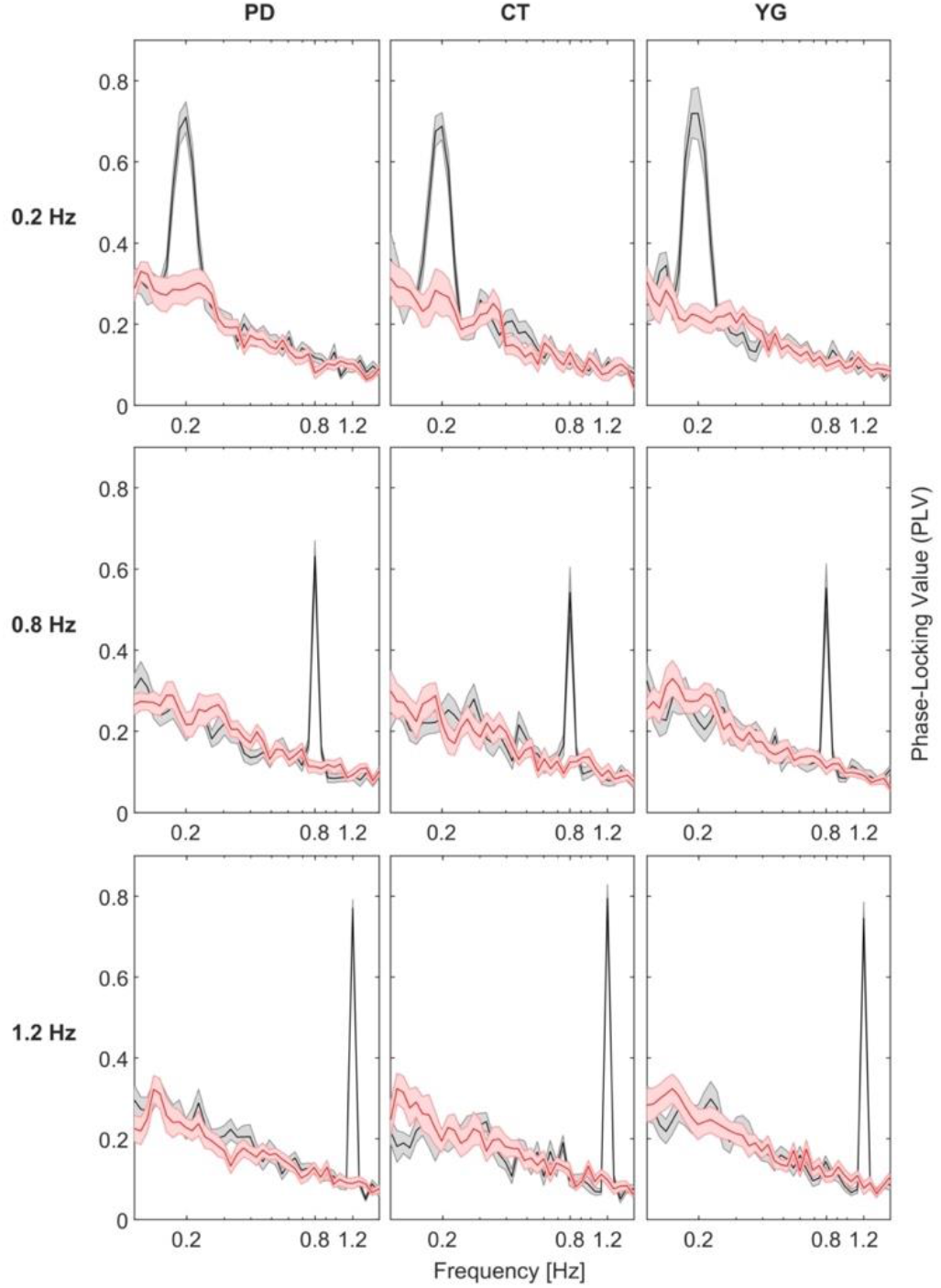
Global phase-coupling spectra of COP in A-P direction. Rows show condition (frequency of the tunnel motion), columns show groups of participants. Black solid lines indicate mean across subjects within each group, with grey shaded areas representing SEM. Analogously, red graphs represent analysis of the baseline (static tunnel) with the respective wavelet parameters at each condition.

**Fig. 5.**
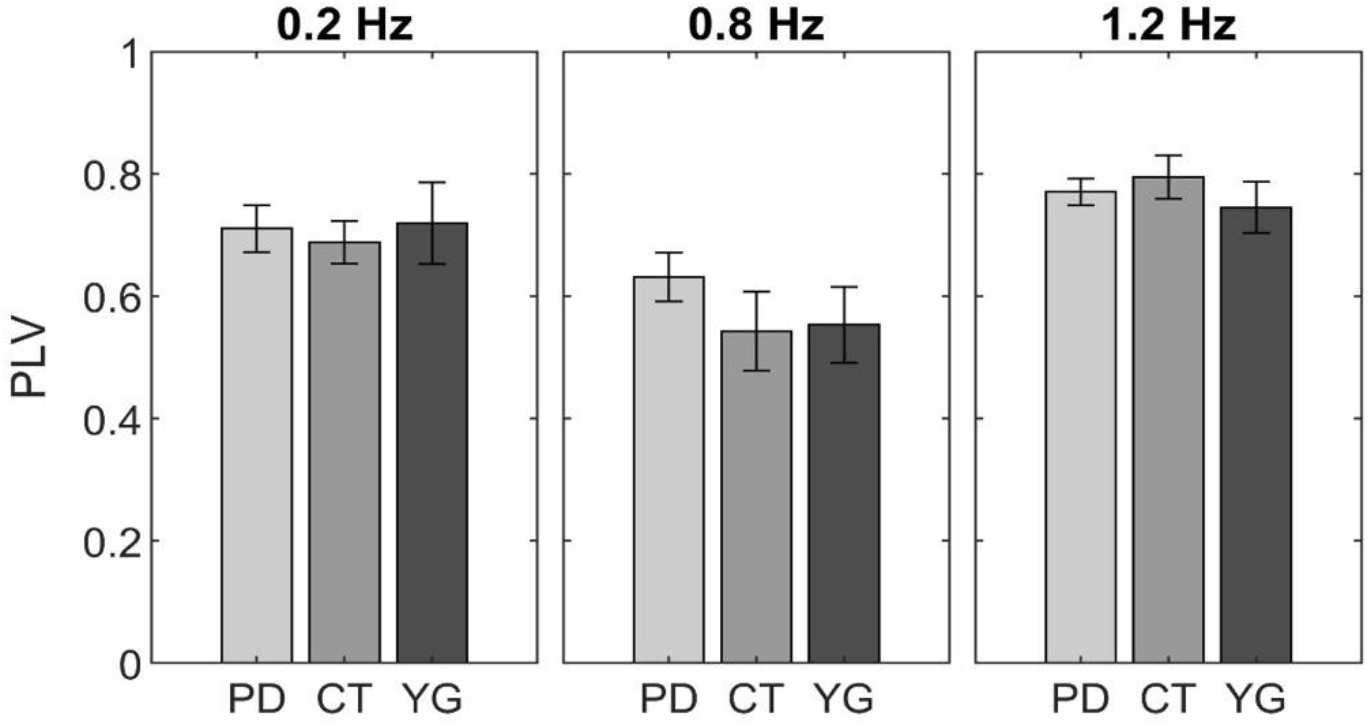
Average PLV magnitude at each stimulus frequency obtained from the COP spectra. Height of the bars displays absolute PLV values at each respective frequency band of stimulation, error bars indicate SEM across participants within each group. Columns represent stimulus condition, different grey shadings represent groups (PD, CT, YG).

**Fig. 6.**
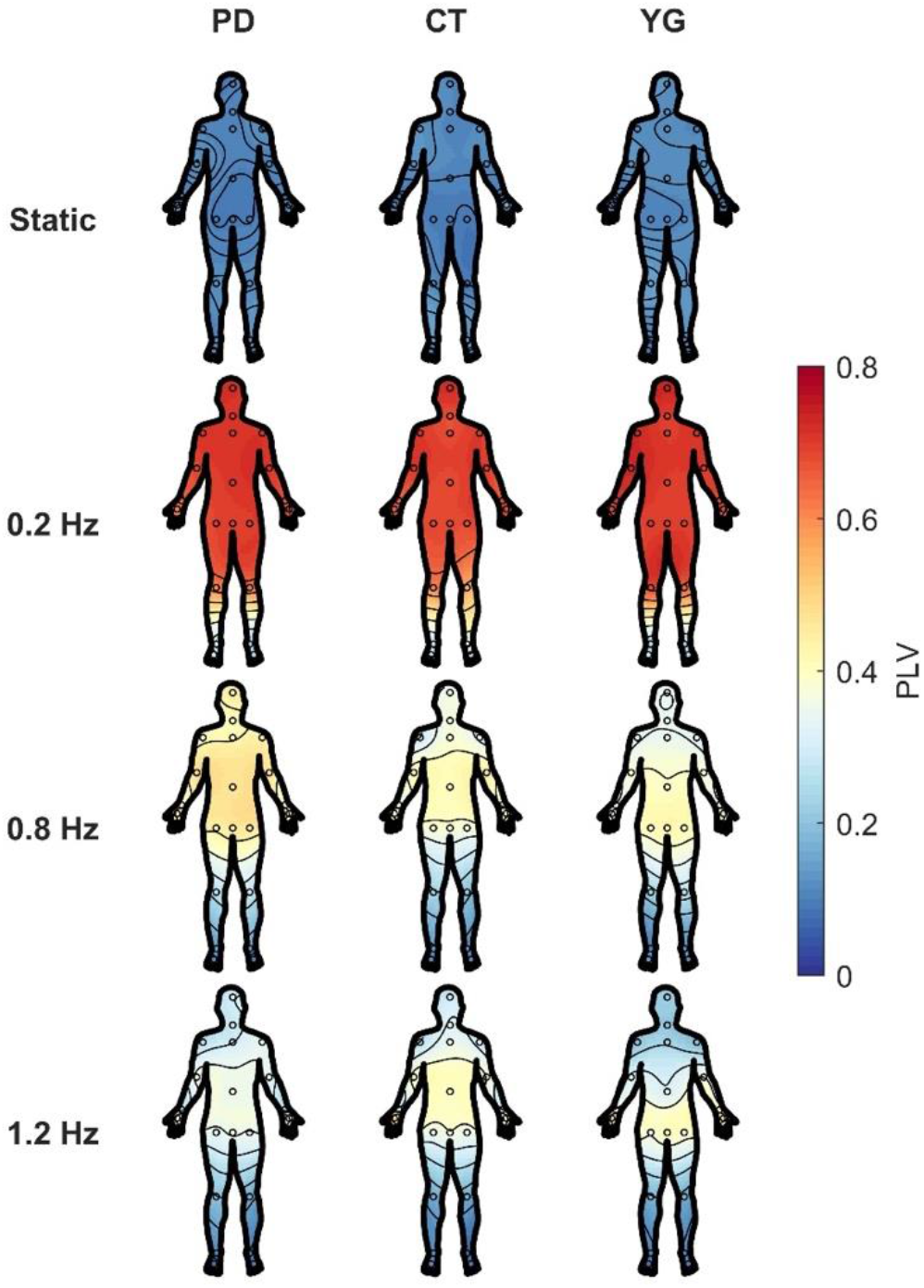
PLV of the 25 body segments obtained by the Kinect in A-P direction. Color-coding represents group average of PLV for each body segment at each respective frequency band of stimulation. The circles represent single body segments. Columns from left to right display groups (PD, CT, YG). Rows from top to bottom show different movement conditions of the tunnel (Static, 0.2 Hz, 0.8 Hz, 1.2 Hz).

PLV magnitude of the body segments at the respective frequency band for each stimulus are visualized in Figure 6. For stimulation at 0.2 Hz, all three groups showed a strong phase coupling to the visual stimulation across their whole body. For stimulation at 0.8 Hz, overall phase-locking became weaker. Here, PD patients showed homogeneous coupling to visual stimulation from their hip upwards. The elderly control group exhibited comparably weaker phase-locking around the head and shoulders, with phase-locking concentrating around the torso. This effect was even stronger in the group of young healthy adults. At visual stimulation of 1.2 Hz, PD patients only showed weak coupling, albeit still rather homogeneously distributed across their upper body. Here, the concentration of phase-locking around the torso for the healthy elderly group shifted downwards towards the hip. At the highest frequency, the young group exclusively phase-locked to the visual stimulus with their hips. For visualization of the static condition (baseline), analysis with wavelet parameters for the middle frequency of 0.8 Hz was chosen representatively. Responses to the static tunnel at the other frequency bands were analogous. As we wanted to put emphasis on how PLV distributed across body segments rather than single PLV magnitudes, we calculated a shoulder-to-hip ratio (SHR). For this purpose, we first normalized PLVs across the body for each subject for better comparability. We then obtained the average PLV of the three segments representing the hip as well as of the three segments representing the upper torso around the shoulders. Subsequently, we calculated the difference in phase-locking between upper and lower torso by subtracting the average of the shoulder segments from the average of the hip segments. Positive values correspond to stronger phase-locking around the hip, negative values correspond to stronger phase-locking around the shoulders. A value of 0 would correspond to identical phase-locking across the torso. SHR for all groups at each motion condition is displayed in Figure 7.

**Fig. 7.**
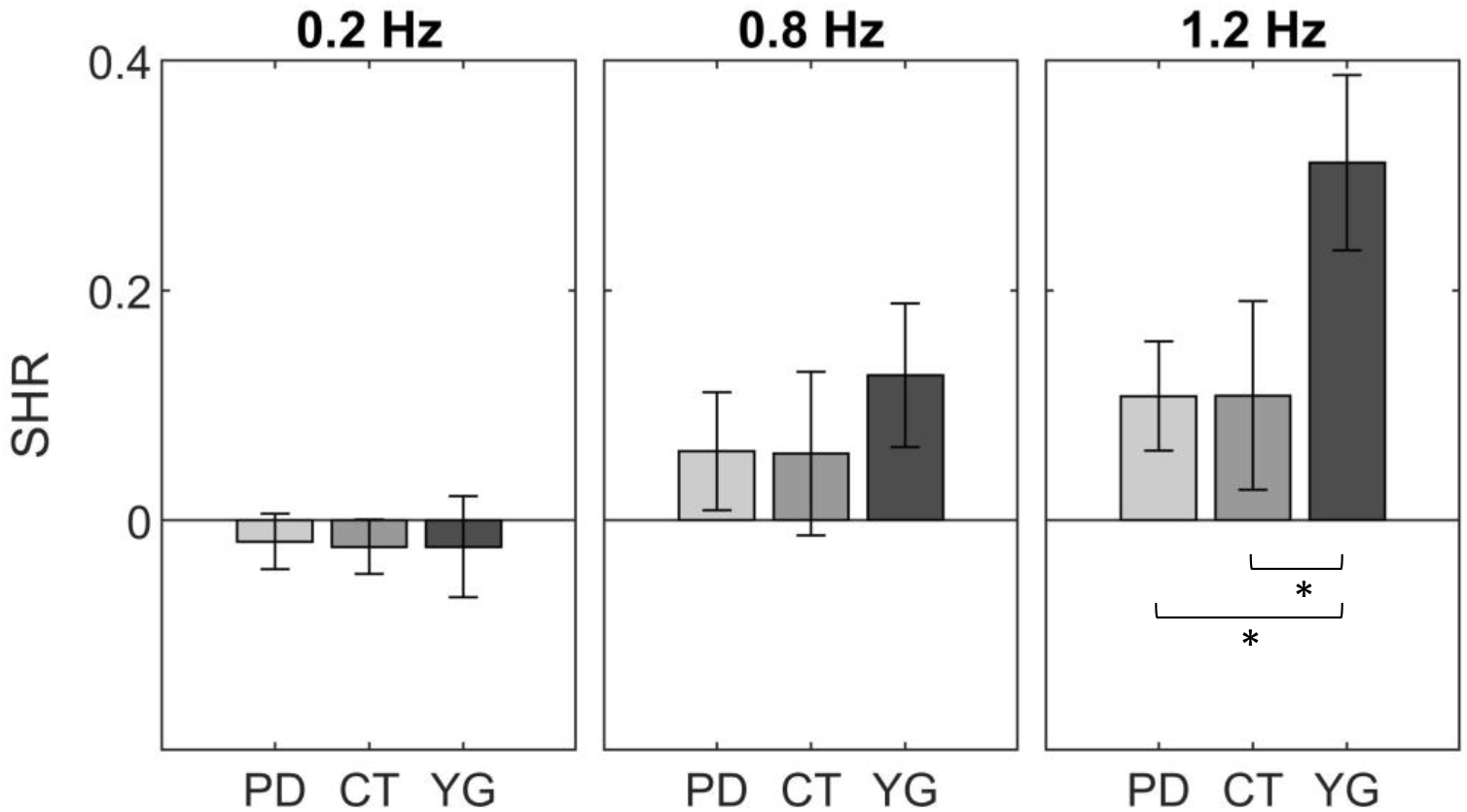
Average shoulder-hip-ratio (SHR, difference in PLV between hip and shoulders) at each stimulus frequency obtained from body segment PLVs. Magnitude of the bars displays SHR at each respective frequency band of stimulation, error bars indicate SEM across participants within each group. Columns represent stimulus condition, different grey shadings represent groups (PD, CT, YG).

For stimulation at 0.2 Hz, all groups showed small negative SHR, indicating that their shoulders phase-locked to the stimulus slightly stronger than the hips. The difference between hip and shoulders was almost identical across groups (H(2)=.76, p=.685). For stimulation at 0.8 Hz, average phase-locking of the hip was stronger than phase-locking of the upper torso, indicated by the positive difference values. Here, there was also no significant difference in SHR between groups (H(2)=1.16, p=.561). Stimulation at the highest frequency of 1.2 Hz revealed positive SHR (hips show stronger phase-locking than shoulders) for all groups. At this highest frequency, however, there was a significant effect of the group (H(2)=7.67, p=.022). Here, the young healthy participants showed significantly more phase-locking in the hips than in their shoulders in comparison with the other groups (YG-PD: p=.036, r=.44; YG-CT: p=.074, r=.43). Noteworthy, there was no significant difference between patients and age-matched control subjects (PD-CT: p= 1.000, r=.01). The frequency of the tunnel motion had no significant effect on the SHR of the PD patients (χ^2^(2)=4.44, p=.115) and of the age-matched controls (χ^2^(2)=3.50, p=.174). However, there was a significant effect of the frequency of the tunnel on the distribution of PLV across the body for the young healthy adults (χ^2^(2)=11.20, p=.004). Here, pairwise comparison revealed a significant difference between the lowest frequency of 0.2 Hz and the highest frequency of 1.2 Hz (p=.003, r=.85).

## Discussion

The aim of this study was to investigate multi-segment phase-locking to an oscillatory visual stimulus in individuals suffering from PD and two groups of healthy adults, an age-matched group to the patients and a group of young adults.

Confirming our first hypothesis, analysis of mean sway amplitude revealed PD patients under dopaminergic “on” medication to have significantly larger sway in A-P direction than both the age-matched controls and the young healthy adults. This applied to both COP (Figure 2) and the body segments obtained by a video-based motion capture system (Figure 3) and corroborates previous research (Doná et al., 2016; Cruz et al., 2018). The qualitative distribution of MSA across the body reflected a strongly pronounced ankle strategy (Winter et al., 1998; Peterka, 2002) in the group of PD patients, with largest sway amplitudes around the head which decreased towards the lower body. Distribution was similar in both other groups, but with equally weaker MSA values. Increased head sway of PD patients, indicating a more pronounced ankle strategy, has previously been reported by Chastan and colleagues (2008) and might reflect increased rigidity of the body which often accompanies PD. Remarkably, motion of the tunnel had no significant effect on overall MSA for PD, but small effects on both CT and YG. The generally higher sway of PD patients as compared to the other groups was thus not affected by the visual condition. This is in line with results from related studies (Cruz et al., 2018, 2021).

Global phase-locking spectra (Figure 4) revealed a highly consistent phase of the COP signals in A-P direction at each frequency of stimulation, for all tested groups. This indicated a strong response to the visual stimuli, as all participants phase-locked their COP only at the frequency at which the tunnel oscillated. Statistical analyses confirmed, with only one exception at 1.2 Hz between PD and CT, that the magnitude of PLV at each condition was not significantly different between groups. In addition, the small standard errors indicated by the shaded areas in Figure 4 and the error bars in Figure 5 reflect little deviation and thus stable responses within each group. This confirms phase-locking to be a strong and stable effect of periodical visual stimulation on body sway (Engel et al., 2021). PLV was significantly affected by frequency within each group. In each case, however, phase-locking only differed significantly in pairwise comparison of the lowest frequency of 0.2 Hz to the higher frequencies of 0.8 Hz and 1.2 Hz. PLVs were lowest for the 0.8 Hz condition and highest for the 1.2 Hz condition in all groups. This behaviour has been observed in healthy adults before (Engel et al., 2021) and might be explained by a mode switching process to adapt to different frequencies (Creath et al., 2005), for which 0.8 Hz indicates a transitional state. This idea, however, would need to be tested by evaluating additional frequencies in the range between 0.2 and 1.2 Hz.

Considering these first results regarding phase-locking of participants’ COP, we must reject our second hypothesis, since there was no difference in phase-locking to the stimulus between PD patients and age-matched controls despite their different sway magnitudes. Moreover, young adults were affected by and phase-locked to the visual perturbation in the same way as older healthy adults and PD patients at all frequencies. Similar time-frequency responses of COP to visual stimuli in young and elderly healthy adults have also been found by Loughlin & Redfern (2001). This suggests that the phase-locking of body sway reflected in COP-PLV is neither affected by PD nor by age, independent of frequency power and sway magnitude.

Overall, PLV analysis of COP signals revealed no effect of visual stimulation on sway magnitude but did reveal an effect on PLV. In addition, sway magnitudes that were significantly different between groups stood in contrast with equally strong PLV across groups. This means that during visual perturbation, the initial amplitude of body sway and thus stability was maintained, and each perturbation was successfully counteracted by the sensorimotor system phase-locking the body sway to the visual motion in the respective direction. This maintenance of body sway amplitudes was successfully achieved by both groups of healthy adults and PD patients alike. By evaluating PLV of the COP, we were able to show that PD patients can compensate for periodic visual perturbation to maintain their balance. While applying a new method to evaluate phase responses to the stimuli, we were hence able to confirm related findings (Cruz et al., 2018, 2021). These findings suggest that, while PD patients exhibit impaired motor output, their sensorimotor processing of visual information remains intact. Potentially, this result reflects that patients in our study were in rather early stages of the disease; it remains an open question if this ability to compensate for visual perturbation is maintained with disease progression.

Incorporating additional full body motion tracking allowed us to test our third hypothesis, that PD patients use a different strategy to compensate for the visual perturbation. This was evaluated by PLVs of the 25 body segments recorded by a motion-capture system for each group and frequency (Figure 6). With increasing frequency of the tunnel, PD patients maintained a rather homogeneous distribution of PLV across their body, while phase-locking of the age-matched control subjects centred slightly towards the lower torso. The group of young healthy adults exhibited the most prominent shift of PLV distribution with increasing frequency, as towards the highest frequency, they phase-locked exclusively with their hip. This finding is remarkable for several reasons. Since there was no difference in PLV of the COP between groups, this indicated an equal ability to adapt to the visual motion. However, when looking at different body segments, it became apparent that the groups used different strategies to achieve this task. To maintain balance, it is crucial to keep the body’s centre of mass within a certain area and range of sway (Horak & MacPherson, 1996; Peterka, 2002; Horak, 2006). Stabilizing the centre of mass, especially at higher frequencies, was thus achieved through different strategies by each group, which was neither observable by sole investigation of COP nor by sway magnitude across the body. Moreover, the prominent shift in strategy of the young group has been observed in healthy adults before (Engel et al., 2021). As a result of the now visible difference in PLV distribution across the body between groups, especially at the torso, we quantified the ratio of phase-locking across the torso by comparing PLVs between the hip and the shoulders (Figure 7). This confirmed the young participants to have significantly more phase-locking around their hip at the highest frequency of 1.2 Hz, with no significant differences between patients and age-matched controls. This could mean that young healthy adults shifted their ankle strategy towards a hip strategy as frequency of the perturbation increased (Nashner et al., 1989; Winter, 1995; Boehm et al., 2019). They were able to adjust to the visual perturbation by switching modes within their body sway (Creath et al., 2005). Even though they were still able to compensate for the perturbations, PD patients and age-matched control subjects maintained their ankle strategy to a much larger degree as frequencies increased. This might be explained by a larger rigidity of the overall body in both groups. Thus, PD patients indeed exhibited more rigidity and less ability to adapt their postural strategy while compensating for balance perturbations (Bloem, 1992; Horak et al.,1992; Chong et al., 2000; Schoneburg et al., 2013). This confirms our third hypothesis. However, the same holds true for age-matched controls, which might indicate an age-effect, rather than an effect of the disease (Hsu et al., 2013).

The larger sway in regard to amplitude but apparently unaffected phase-coupling of the PD patients, even when compared with the group of young healthy adults, could have two explanations. First, it has been proposed that movements elicited by external cues, as was the case in our study, might bypass the basal ganglia and therefore are less impaired in PD patients (Cunnington et al., 1995). Second, the unimpaired responses might be linked to the L-dopa medication which all our participants received at the time of testing. There is evidence that dopaminergic medication does not improve neuromuscular postural stability (Koller et al., 1989; Bloem, 1992; Grimbergen et al., 2009) and effects of L-dopa on other systems contributing to postural responses are under debate (Feller et al., 2019). For instance, Chen et al. (2016) concluded from their study that L-dopa medication does not improve stability of the neuromuscular system but improves responsiveness of the sensorimotor system, which is strongly supported by our findings.

Limitations of our study included the relatively small and unequal groups of participants, accompanied by a rather large variety of age, disease progression and equivalent dose of L-dopa in the group of PD patients. Given the large variability within PD itself regarding age of onset and disease progression rates, on the other hand, our patients sample represents a rather typical cohort of PD patients in early and mid-disease stages. In addition, we were unable to exactly match the distribution of sex between patients and both control groups. As for the stimulus, we only applied three different frequencies, which was mainly to prevent fatigue in the patients and elderly control subjects, as we aimed to obtain 10 trials per condition with sufficient stimulus duration.

Considering the newly observed shift in strategy towards the hip for higher frequencies of visual perturbation, in which the young group significantly differed from the control subjects and patients, it would be intriguing to conduct experiments with larger groups as well as adding smaller frequency increments up towards higher frequencies of visual stimulation. This would allow possible observations on how and where in the spectrum strategy shifts occur. Moreover, a larger clinical population would allow investigation of subgroups to gain insight into how these strategy shifts are affected by factors like disease severity and medication. Our phase-locking analyses in combination with full-body motion tracking provided new insight into postural responses to visual perturbations. Combined with the mobility of our setup and its potential to expand the reach of experiments independent of a research laboratory, this newly introduced technique seems promising for further applications, especially in regard to biomarkers in different clinical populations where balance and postural control are impaired.

## Conclusion

Applying a low-cost and mobile setup to visually perturb balance in a clinical population of patients suffering from PD, as well as an age-matched control group and a group of young healthy adults, we successfully evaluated three hypotheses: Firstly, we found PD patients to have significantly more body sway regarding sway magnitude, irrespective of visual stimulation. Secondly, PD patients were able to phase-lock to and thus compensate for the visual perturbations in the same way as age-matched control subjects and young healthy adults. For PD, these two findings indicate impaired neuromuscular stability but intact visuomotor control. Thirdly, as frequency of the stimulation increased, we found significant differences in postural strategies between the groups, which have previously not been reported in this manner. This was achieved by a combination of phase-locking analyses and full-body motion tracking. Our newly introduced technique revealed aspects of postural responses to perturbations that investigations of sway magnitude and frequency power could not provide, and which might enrich research on postural control for future applications in various settings.

## Acknowledgments

Research was funded by Deutsche Forschungsgemeinschaft (DFG): IRTG-1901, CRC/TRR-135 (project number 222641018) and EU: PLATYPUS.

